# Analysis of new nosological models from disease similarities using clustering

**DOI:** 10.1101/2020.04.10.035394

**Authors:** Lucía Prieto Santamaría, Eduardo P. García del Valle, Gerardo Lagunes García, Massimiliano Zanin, Alejandro Rodríguez González, Ernestina Menasalvas Ruiz, Yuliana Pérez Gallardo, Gandhi Samuel Hernández Chan

## Abstract

While classical disease nosology is based on phenotypical characteristics, the increasing availability of biological and molecular data is providing new understanding of diseases and their underlying relationships, that could lead to a more comprehensive paradigm for modern medicine. In the present work, similarities between diseases are used to study the generation of new possible disease nosologic models that include both phenotypical and biological information. To this aim, disease similarity is measured in terms of disease feature vectors, that stood for genes, proteins, metabolic pathways and PPIs in the case of biological similarity, and for symptoms in the case of phenotypical similarity. An improvement in similarity computation is proposed, considering weighted instead of Booleans feature vectors. Unsupervised learning methods were applied to these data, specifically, density-based DBSCAN clustering algorithm. As evaluation metric *silhouette* coefficient was chosen, even though the number of clusters and the number of outliers were also considered. As a results validation, a comparison with randomly distributed data was performed. Results suggest that weighted biological similarities based on proteins, and computed according to cosine index, may provide a good starting point to rearrange disease taxonomy and nosology.

## I. Introduction

Classical human disease classification has traditionally been focused on empirical phenotypical features such as symptoms, anatomy and physiology, rather than on molecular etiology [1]. In the last decades, advances in molecular biology have allowed science to describe human disease in terms of molecular and genetic characteristics [2], thanks to interrelated large amounts of very different types of data [3]. It is known that incorporating these advances about disease’s molecular and genomic variations in their diagnostic and treatment could transform drug development and medicine [4].

Some studies agree that in order to organize the current disease knowledge, a categorization capable of representing disease taxonomy should be performed. Categorization is by nature a holistic process, demanding a global picture of the organization and mechanistic details of individual components. In the human disease case, this categorization from a global point of view could come with a better understanding of the cause and effects of diseases [3]. However, many common human diseases are still diagnosed as if they were homogeneous entities, using criteria that have hardly evolved in more than a century [4]. Therefore, it is necessary to research on the development of new disease taxonomies and classification systems that are more suitable to include molecular information, combining these molecular and biological data with physiological and phenotypical data [5].

The present works intends to research on the possibilities to create new nosologic methods of disease classification from different types of similarities based on biological and phenotypical traits. A literature review of the disease classification systems and the computational approaches to generate them was performed. The methods include the f data acquisition from DISNET platform and similarity coefficients computation processes. Unsupervised learning techniques were applied to categorize diseases and then validated and evaluated.

## II. RELATED WORK

Classification has evolved over the years: from Linneo, who in 1763 classified diseases as *exanthematics, phlogistics* and *dolorous* [6], to Wilbur’s Manual of the International List of Causes of Death [7], which in 1909 still lacked of distinction between nowadays differentiated diseases, such as type I and type II diabetes. Currently, the main classification systems derive from the different vocabularies used in medical literature and research, including but not only International Classification of Diseases (ICD), Medical Subject Headings (MeSH), Systematized Nomenclature of Medicine – Clinical Terms (SNOMED-CT), Unified Medical Language System (UMLS) and Disease Ontology (DO).

Human disease classification nowadays relies in the observational correlation between pathologic analysis and clinical syndromes. Disease characterization in such way has stablished a useful nosology for physicians, defining nosology as the branch of medicine dedicated to disease description and classification. However, it has significant limitations regarding modern medicine that reflect both a lack of sensitivity when identifying preclinical disease states, and a lack of specificity in defining diseases unequivocally. A human disease classification combining conventional reductionism with systems biomedicine non-reductional approach is required in order to include the high volume and heterogeneous genomic, proteomic, transcriptomic and metabolic data not taken into account thus far [8].

In 2011, a call to reform disease taxonomy in order to promote the inclusion of the last scientific advances was made [4]. At that time, the USA National Academy of Sciences (NAS) formed a committee to analyze the feasibility and necessity of a “new taxonomy of the human disease based in molecular biology”, as well as a model framework to develop it [5]. As disease taxonomy plays a fundamental role defining diagnosis, treatments and mechanisms of human disease [9], a new taxonomy reflecting the characteristics proposed by the NAS would greatly facilitate *precision medicine* development.

Classifying diseases into groups determines the response that should be applied in presence of a disease classified within a certain group [10]. If this classification is modernized incorporating the known or inferred disease molecular information, the classification would not only provide the classical structure built on disease physiology, but also provide insight about the associations between disease groups to specific diagnostics and treatments [8], [11].

The taxonomies developed so far still reflect the lack of inclusion of molecular information when classifying diseases, but are a good starting point or object of comparison to new taxonomies in the absence of a gold standard to evaluate [12]. Works [11] and [9] are a proof of this. In [11], new diseases hierarchies were inferred by a new method called *ParentPromotion* using disease – gene data and comparing them to both MeSH category C forest and DO subnetworks. It was proved that part of MeSH and DO hierarchical structures could be deduced from molecular data. The algorithm *ParentPromotion*, based on hierarchical clustering, converted the generated difficult-to-interpret dendrogram into a disease hierarchy based in the number of PubMed publications related to each disease. In [9], a *New Classification of Diseases* (NCD) was proposed developing an algorithm that predicted additional categories of a disease by integrating multiple networks consisting of disease phenotypes and their molecular profiles. An improvement in the current code structure of ICD (concretely of the version ICD-9-CM, *International Classification of Diseases, 9th Revision, Clinical Modification*) was proposed by demonstrating that a substantial number of diseases (cancer and infectious diseases among others) tended to have molecular connections with diseases in other chapters. The new proposed classification integrated disease networks taking into account molecular and phenotypic connectivity among diseases and predicted new disease classes.

One of the approaches that can be addressed to research in the development of new taxonomies is the *Network Medicine* [13]. In it, not only the molecular complexity of a particular disease is systematically explored, but also the molecular relationships among apparently distinct pathophenotypes [14]. To this end, Human Disease Networks (HDN) are built. HDN is a global concept that refers to a complex network in which its nodes are the diseases or disorders and its links, the relationships between them. Such relationships can be stablished based on the sharedness of multiple traits, being two of the most important the disease genes [15] or the associated symptoms [16].

Moreover, the use of semantic similarity between biological processes to estimate disease similarity could enhance the identification and characterization of disease similarity [17]. In 2011, Li *et al*. computed semantic similarity according to ten different measures based on DO ontological terms shared between diseases [18]. In 2014, a new method called *SemFunSim* to measure disease similarity by integrating semantic and gene functional association was developed [19]. The same year, Sun *et al*. were able to predict disease associations by analyzing a biological network in three ways: annotation-based, function-based and topology-based [20]. In 2015, Kim *et al*. proposed a method that consisted in the construction of disease-gene and disease-drug literature relationships matrices, from which similarities were calculated based on the mutual information [21]. In 2016, DisSim was introduced, an online system to explore significant similar diseases and exhibiting potential therapeutic drugs, using different algorithms. Carson et al. built, in 2017, a disease similarity matrix based on the uniqueness of shared genes using OMIM (Online Mendelian Inheritance in Man) and DO annotations [22].

## III. METHODS

### A. DISNET project and data acquisition

The current work has been developed in the context of DISNET system [23], a web service to extract disease knowledge structured in the basic concepts of the *Human Disease Networks* (HDN) [15], [16]. Although at this moment only its phenotypical data is publicly available, the system aims to form a complex multilayer graph. In the present, it has a two levels topology: the phenotypical layer and the genetic or biological layer. The first layer is based on the extraction and storage of phenotypical disease knowledge, mainly disease symptoms. This information is obtained by literature mining processes of disease descriptions from different sources (such as Wikipedia^1^ or Pubmed^2^). The second layer is supported on the inclusion of genetic and/or biological disease knowledge. In particular, information about genes, proteins, metabolic pathways and Protein-Protein Interactions (PPIs) related to diseases is extracted from DisGeNET ^3^, WikiPathways ^4^ and IntAct ^5^. This way, the final aim of DISNET is the construction of the total network where relationships between diseases can be analyzed from different points of view, providing a greater understanding of diseases. Moreover, such approach could help finding out not only the phenotypical and genetic features shared by diseases, but also which drugs have a high probability of being reused for new indications, meeting a drug repurposing sense.

### B. Disease similarity coefficients computation

The disease similarity computation process is underpinned by Vector Space Model (VSM) methods [24], in which diseases are represented as vectors of features. In the present work, those features are phenotypical (namely, disease symptoms) and genetic or biological traits (namely, disease genes, proteins, pathways and PPIs). Vector operations can be used to compare diseases and quantitatively illustrate the similarity between them in terms of each trait. In this context, different measures have been used through the literature to compute disease similarity. We chose three metrics as suggested by [25]: cosine index, Jaccard index and Dice index. Such metrics are commonly used to compute semantic similarity between documents defined as Boolean vectors of features. Translating this to our approach, A and B would be two vectors related to two different diseases representing whether a biological or phenotypical trait is associated (1) or not (0) to their respective disease. Having a Disease – Gene Association (DGA) score available made possible going one step further in the disease similarity computation by weighting some of the biological features’ vectors. The DGA score was obtained from DisGeNET [26] and enabled building vectors of real numbers of genes, proteins and metabolic pathways. The weighted disease similarity was computed as follows:

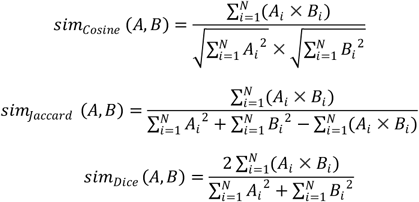

where A and B are two features vectors of real numbers related to two different diseases. In the case of proteins and pathways, to weight the vectors, it was neccesary to go through the genes. A gene could encode more than a protein and may be involved in multiple metabolic pathways (also, a metabolic pathway will have several genes involved in it). To weight protein and pathway vectors, the largest DGA score of the genes related to these features was considered.

Combining the different indexes and features, there are 24 similarity coefficients, of which 15 comes from Boolean features vectors and 9 from weighted features vectors. For the Boolean vectors, the features are genes, proteins, metabolic pathways, PPIs and symptoms, and for the weighted vectors, the features are genes, proteins and pathways. Each type of features vector combined with each of the three indexes (cosine, Jaccard and Dice) result in the named 24 similarity coefficients.

### C. Clustering expermients

The aforementioned 24 similarity coefficients were organized in similarity matrices in which 3,671 diseases were combined two-by-two without repetition. The 3,671 diseases were both present in the phenotypical and in the biological layer. One similarity matrix for each different similarity coefficient was generated, in which there were as many columns and rows as diseases. The value in each field of the matrix corresponded to the similarity between the disease of the column and the disease of the row. Therefore, these square matrices were symmetric and all the elements in the diagonal were equal to “1”. Before applying clustering techniques to the data, matrices similarities would be converted to distances so that *distance = 1 – similarity*.

Once obtained the 24 different distance matrices, clustering analysis was performed. Unsupervised learning can facilitate the task of forming new diseases groups under the lines of modernizing the current classification systems. According to the data we held and since the number of diseases groups to form remained unknown, density-based clustering methods were chosen. In particular, DBSCAN (*Density-Based Spatial Clustering of Applications with Noise*) [27] was used. DBSCAN is designed to find core samples of high density and expands clusters from them. It requires two parameters: *Eps* and *MinPts*. On the one hand, *Eps* (ε) is the parameter used to specify the radius of a neighborhood considered for every object, that is to say, the maximum distance between two samples for one to be considered as in the neighborhood of the other. *MinPts* is the parameter that determines whether a neighborhood is dense or not, it specifies the density threshold of dense regions.

For each distance coefficient, several *Eps* and *MinPts* were used in order to explore the different results obtained. *Eps* took the values 0.3, 0.4, 0.5, 0.6, 0.65, 0.7, 0.75, 0.8, 0.85, 0.9, 0.95 and 0.99, while *MinPts* was varied between 2, 3, 5 and 30.

### D. Evaluation

Due to the purpose of this work of obtaining new groups of diseases, there was not an available ground truth which clustering results could be compared to. The aim was not to generate models that predicted disease groups in the already existing classification systems but providing novel categorizations. Therefore, it was necessary to use an intrinsic method to assess the clustering quality, which could examine the heterogeneity between the objects of the different clusters and the homogeneity among the objects within a cluster [28].

One of the most popular index used in the literature as an internal clustering evaluation method is *silhouette* coefficient, which measures the cohesion and separation of the generated clusters [29]. *Silhouette* ranges from −1 to +1, where values close to +1 indicate that the objects are well matched to their own cluster and poorly matched to neighbor clusters, and values of −1 indicate that the clustering configuration may have too many or too few clusters. *Silhouette* coefficient was the chosen metric to evaluate the obtained disease groups, considering it as a guidance of how well the diseases are distributed along the clusters.

Another parameter that was considered when determining the quality of a clustering result, was the number of outliers, namely those points that were not grouped within any cluster. If most diseases are not inside a cluster, then the clustering model will not be suitable to generate a new disease nosology. On the contrary, a result in which all the data points were grouped as just one did not provide any nosologic information and had a low quality for the current task. Therefore, when determining the best results, both *silhouette* coefficient, the number of clusters formed, and the number of outliers were taken into account.

Furthermore, in the cases of the results with higher silhouette, another validation was performed. Such results were compared to the same experiments but on 100 simulations shuffling elements by columns in the features arrays. That is, the distributions of a feature along diseases was the same but the similarity matrices were randomized, ensuring triangle inequality… Besides the 24 distances matrices, DBSCAN was implemented on the distance matrices generated from different probability distributions: uniform, beta, gamma and normal distributions. This way, it could be proven that those experiments obtaining the highest quality were also better than the results when performing DBSCAN on the data generated from different random probability distributions.

## IV. RESULTS AND DISCUSSION

All results can be found at a public repository^6^, where both code and results are attached. From all the possible combinations between the values of *Eps* (0,3; 0,4; 0,5; 0,6; 0,65; 0,7; 0,75; 0,8; 0,85; 0,9; 0,95 and 0,99) and *MinPts* (2, 3, 5 and 30), those leading to a higher *silhouette* coefficient are showed here. They are divided in results obtained by performing DBSCAN on biological similarities (Table I), on phenotypical similarities (Table II) and on weighted biological similarities (Table III). In each table, values of the parameters (*Eps* and *MinPts*) set are shown, as well as obtained *silhouette* coefficient, number of clusters and number of outliers.

**TABLE I.**
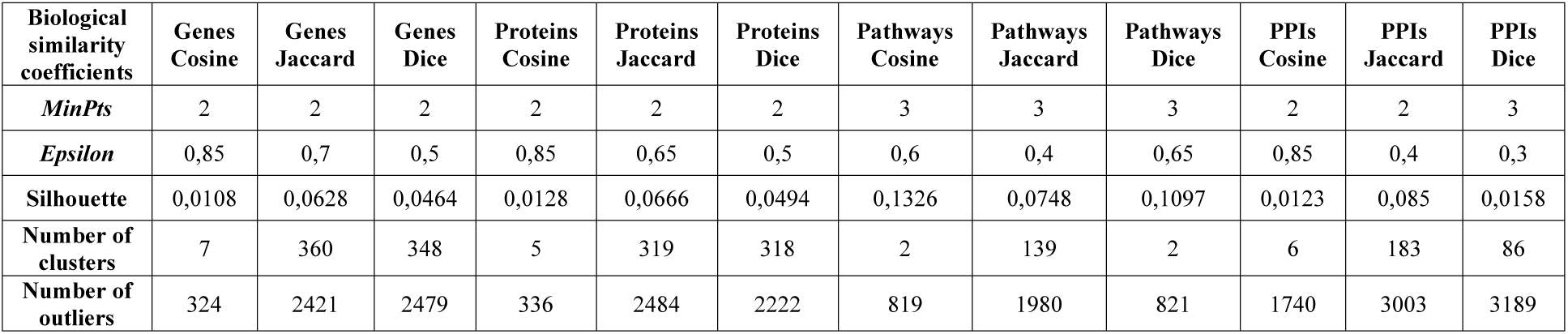
BEST RESULTS FOR BIOLOGICAL SIMILARITIES

| Biological similarity coefficients | Genes Cosine | Genes Jaccard | Genes Dice | Proteins Cosine | Proteins Jaccard | Proteins Dice | Pathways Cosine | Pathways Jaccard | Pathways Dice | PPIs Cosine | PPIs Jaccard | PPIs Dice |
| --- | --- | --- | --- | --- | --- | --- | --- | --- | --- | --- | --- | --- |
| *MinPts* | 2 | 2 | 2 | 2 | 2 | 2 | 3 | 3 | 3 | 2 | 2 | 3 |
| *Epsilon* | 0,85 | 0,7 | 0,5 | 0,85 | 0,65 | 0,5 | 0,6 | 0,4 | 0,65 | 0,85 | 0,4 | 0,3 |
| *Silhouette* | 0,0108 | 0,0628 | 0,0464 | 0,0128 | 0,0666 | 0,0494 | 0,1326 | 0,0748 | 0,1097 | 0,0123 | 0,085 | 0,0158 |
| Number of clusters | 7 | 360 | 348 | 5 | 319 | 318 | 2 | 139 | 2 | 6 | 183 | 86 |
| Number of outliers | 324 | 2421 | 2479 | 336 | 2484 | 2222 | 819 | 1980 | 821 | 1740 | 3003 | 3189 |

**TABLE II.** BEST RESULTS FOR WEIGHTED BIOLOGICAL SIMILARITIES

| Weighted biological similarity coefficients | Genes Cosine | Genes Jaccard | Genes Dice | Proteins Cosine | Proteins Jaccard | Proteins Dice | Pathways Cosine | Pathways Jaccard | Pathways Dice |
| --- | --- | --- | --- | --- | --- | --- | --- | --- | --- |
| *MinPts* | 2 | 2 | 2 | 2 | 2 | 2 | 3 | 2 | 2 |
| *Epsilon* | 0,4 | 0,65 | 0,5 | 0,4 | 0,65 | 0,5 | 0,5 | 0,3 | 0,3 |
| *Silhouette* | 0,2013 | 0,1373 | 0,1538 | 0,2217 | 0,1484 | 0,1681 | 0,0681 | 0,0837 | 0,0477 |
| Number of clusters | 463 | 421 | 439 | 460 | 428 | 448 | 3 | 347 | 316 |
| Number of outliers | 1898 | 2282 | 2182 | 1801 | 2183 | 2079 | 848 | 2411 | 1828 |

<sup>6</sup> https://github.com/luciaprietosantamaria/CBMS2020

**TABLE III.** BEST RESULTS FOR PHENOTYPICAL SIMILARITIES

| Phenotypical similarity coefficients | Symptoms Cosine | Symptoms Jaccard | Symptoms Dice |
| --- | --- | --- | --- |
| *MinPts* | 2 | 2 | 2 |
| *Epsilon* | 0,85 | 0,7 | 0,5 |
| *Silhouette* | 0,0108 | 0,0628 | 0,0464 |
| Number of clusters | 7 | 360 | 348 |
| Number of outliers | 324 | 2421 | 2479 |

Among all the experiments performed fixing different parameters, the ones that achieved the best results considering *silhouette* coefficient, the number of clusters and the number of outliers, were those obtained from weighted similarities based on proteins and according to cosine index. In such results, *Eps* and *MinPts* had been set to 0.4 and to 2 respectively. The value of *silhouette* coefficient is 0.22, the number of clusters is 460 and the number of outliers is 1801. Although is still half of the total of the diseases, the number of outliers is significantly lower than in other cases. This means that the model would be able to differentiate more diseases and therefore would entail a better nosology. In this model, the maximum number of diseases per cluster was 173, the minimum was 2, the average was 4,065 and the mode, 2. A visualization of the clustering model is included in Fig. 1, where different colors represent different disease groups.

**Figure 1.**
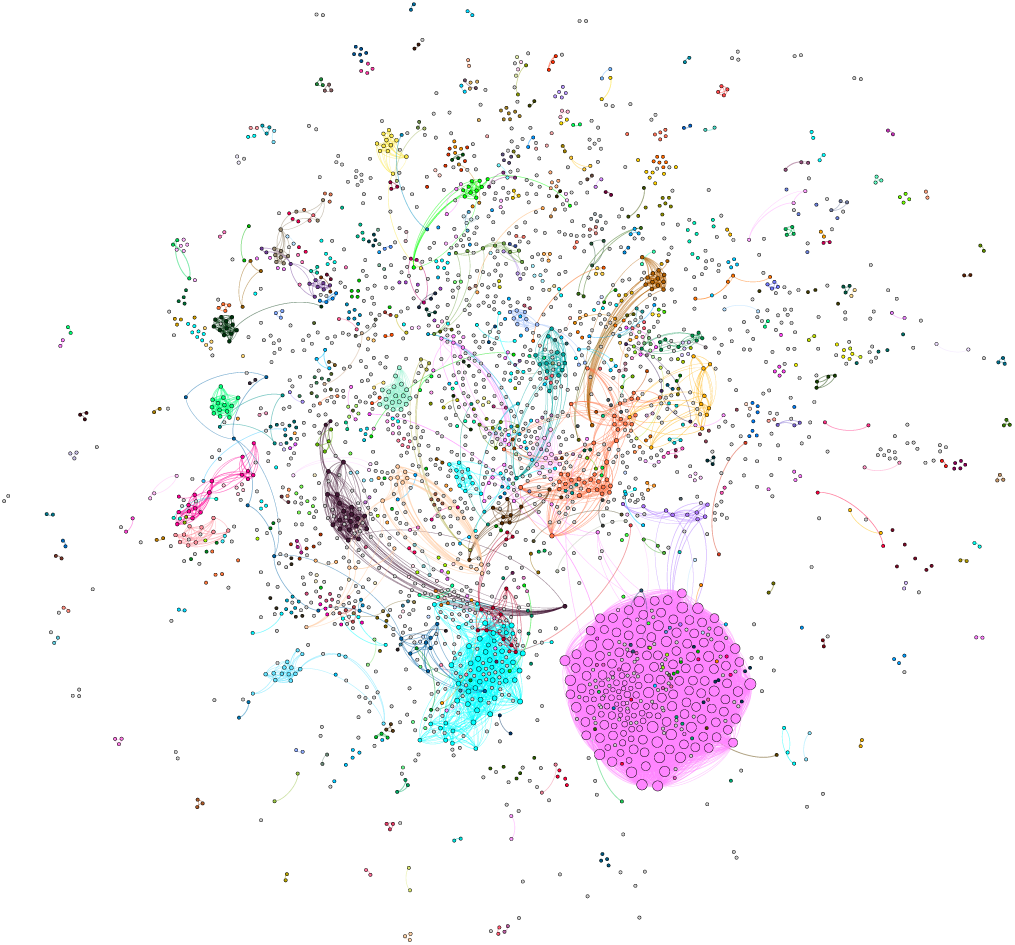
Visual representation of the clusters generated with the weighted protein-cosine similarities.

Performing the same experiment (DBSCAN setting *Eps* to 0.4 and *MinPts* to 2) on analogous data but obtained from randomized similarity matrices, generated the results showed in Table IV. For such randomized matrices, a 100 times simulation was performed, from which we show the mean value ± the standard deviation. In the clustering model for the random similarity matrices, the whole set of points were defined as outliers, with no clusters. Furthermore, and in order to have more insights of the models that would be generated from shuffled feature vectors, DBSCAN setting *Eps* to 0.85 and *MinPts* to 2 was applied to the aforementioned data. *Silhouette* coefficient was negative for all the random distributed experiments (Table V), while actual similarities resulted in positive values of *silhouette*. This led us to think that the similarities that we had computed were meaningful and a good starting point to study disease phenotypical and biological relationships.

**TABLE IV.** COMPARISON BETWEEN RANDOM DISTRIBUTED AND WEIGHTED-PROTEIN SIMILARITY COEFFICIENTS CLUSTERING RESULTS (*EPS* 0.4 AND *MINPTS* 2)

| Similarity coefficients | Shuffled Protein Cosine | Weighted Proteins Cosine | Weighted Proteins Jaccard | Weighted Proteins Dice |
| --- | --- | --- | --- | --- |
| **Silhouette** | $-1 \pm 0$ | 0.22 | 0,10 | 0.14 |
| **Number of clusters** | $0 \pm 0$ | 460 | 267 | 381 |
| **Number of outliers** | $3671 \pm 0$ | 1801 | 2891 | 2460 |

**TABLE V.** COMPARISON BETWEEN RANDOM DISTRIBUTED AND WEIGHTED-PROTEIN SIMILARITY COEFFICIENTS CLUSTERING RESULTS (*EPS* 0.85 AND *MINPTS* 2)

| Similarity coefficients | Shuffled Protein Cosine | Weighted Proteins Cosine | Weighted Proteins Jaccard | Weighted Proteins Dice |
| --- | --- | --- | --- | --- |
| **Silhouette** | $-0,05 \pm 0.001$ | 0.18 | 0,05 | 0.02 |
| **Number of clusters** | $267,71 \pm 17$ | 30 | 216 | 83 |
| **Number of outliers** | $2610,67 \pm 28,45$ | 474 | 1097 | 764 |

Moreover, when contrasting the diseases grouped in different models, some of them agreed in certain clusters. For example, cluster 18 of weighted-genes-cosine similarities model placed exactly the same ten diseases as cluster 20 of weighted-proteins-cosine similarities model, all of them belonging to MeSH class “Cardiovascular Diseases”^7^.

## V. CONCLUSIONS AND FUTURE WORK

A modernization in the current nosology to obtain a better and more comprehensive classification of diseases should be performed in order to include biological and molecular information. This nosology could contribute to improve disease understanding.

The present work has developed an exhaustive analysis of the options for the generation of new nosologic models by applying density-based clustering techniques to data setting different parameters. The used techniques are a state-of-the-art approach. In a future, we will complement the present study with other metrics and methods such as Normalized Mutual Information or K-Means respectively. Some of the conclusions achieved are summarized hereunder.

Disease similarity can be measured as the cosine, Jaccard and Dice index of shared features: genes, proteins, pathways, PPIs and symptoms. When compared with Boolean vectors, weighting feature vectors to compute disease similarity results in more accurate clustering models, with a smaller number of outliers and higher values of *silhouette* evaluation metric.

Density-based clustering algorithms, like DBSCAN, are able to predict groups of diseases letting diseases out of dense regions to be classified as ‘noise’. This is particularly useful for the current work, as not every disease belonged necessarily to a group. A previous analysis of the parameter combination as input to DBSCAN is needed to determine which values of the variables would bring the best result.

Generating new nosologic models of diseases can become an arduous task because of the multiple possibilities. *Silhouette* coefficient can give an insight of which results could be considered as the best from the mathematical point of view. However, silhouette measures the coherence of the grouping and relies on the number of instances. It is not always essential, there are other parameters that should be considered such as the number of outliers or clusters generated.

Regarding the future lines, a new own similarity metric could be proposed. It could be based on a classification system, a mathematical model or a weighting system between the coefficients we already have computed. Moreover, the features vectors could be weighted by other scores and not just by DisGeNET DGA scores. New semantic similarity functions should be tried in order to cover more metrics. Other clustering methods, different from density-based, may be performed. For example, hierarchical clustering could have been interesting to cluster a lower number of diseases set. Important for DISNET system, another future work is to develop a third data layer consisting on the drugs layer, to store disease drugs information.

## Acknowledgment

The work is a result of the project “DISNET (Creation and analysis of disease networks for drug repurposing from heterogeneous data sources applied to rare diseases)”, that is being developed under grant “RTI2018-094576-A-I00” from the Spanish Ministerio de Ciencia, Inovación y Universidades. Lucia Prieto Santamaría’s work is supported by “Programa de fomento de la investigación y la innovación (Doctorados Industriales”) from Comunidad de Madrid (grant IND2019/TIC-17159). Gerardo Lagunes-Garcia’s work is supported by the Mexican Consejo Nacional de Ciencia y Tecnología (CONACYT) (CVU: 340523) under the programme “291114 - BECAS CONACYT AL EXTRANJERO”.

## Footnotes

1 https://www.wikipedia.org

2 https://www.ncbi.nlm.nih.gov/pubmed/

3 https://www.disgenet.org/

4 https://wikipathways.org

5 https://www.ebi.ac.uk/intact/

6 https://github.com/luciaprietosantamaria/CBMS2020

7 https://github.com/luciaprietosantamaria/CBMS2020/blob/master/Results/Cluster18_WGC%20vs%20Cluster20_WPC.txt

